# Biosynthetic diversity and antifungal potential of soil-derived *Saccharomonospora* strains

**DOI:** 10.64898/2026.01.19.700407

**Authors:** Hildah Amutuhaire, Michael Dubovis, Tal Luzzatto-Knaan, Jonathan Friedman, Eddie Cytryn

## Abstract

Soilborne fungal pathogens pose persistent challenges to sustainable agriculture, driving demand for biological alternatives to synthetic fungicides. While actinomycetes, particularly *Streptomyces* have yielded numerous antifungal compounds, less-explored genera, such as *Saccharomonospora* represent untapped sources of novel bioactive metabolites for plant protection. Our previous work identified *Saccharomonospora* as strongly associated with plant disease suppression in organically-amended soils and harbored numerous uncharacterized biosynthetic gene clusters (BGCs). Here, we investigated the biosynthetic capacity and antifungal potential of *Saccharomonospora* using comparative genomics, untargeted metabolomics, and *in vitro* bioassays. Cell-free supernatants from six strains (five soil-derived type strains and one novel isolate) were evaluated against three Fusarium phytopathogens. All strains inhibited at least one pathogen, with *S. xinjiangensis* and *S. viridis* R81 exhibiting the highest and broad-spectrum activity. Metabolomic profiling of the two most bioactive strains and one moderately active strain (*S. cyanea*) revealed that ∼40% of detected metabolites were shared across the three strains although their relative abundances varied. *S. xinjiangensis* and *S. viridis* R81 displayed higher abundances of shared metabolite classes than *S. cyanea*, including alkaloids, polyketides, and peptide derivatives. Comparative genomics across the genus revealed that most BGCs, particularly those encoding non-ribosomal peptides synthetases and polyketides synthases, were strain-specific and had low sequence similarity to characterized BGCs. In contrast, BGCs encoding, indole, ectoine, arylpolyene, and terpene were ubiquitous across the genus. Collectively, these findings demonstrate that *Saccharomonospora* produce antifungal metabolites and harbor diverse, uncharacterized BGCs, positioning this “less-explored” actinomycete genus as a promising source of bioactive compounds for managing soilborne fungal pathogens.

**Importance:** Actinomycetes have historically been rich sources of antifungal metabolites for agriculture and pharmaceuticals, but discovery efforts have focused largely on *Streptomyces*, leaving other genera underexplored. This study demonstrates that *Saccharomonospora*, a rare actinomycete genus, produces antifungal metabolites active against major *Fusarium* pathogens and harbors largely uncharacterized biosynthetic gene clusters, indicating a reservoir of novel chemistry. These findings establish *Saccharomonospora* as a promising yet underutilized resource for discovering new antifungal agents, expanding the toolkit for sustainable plant disease management beyond traditional actinomycete sources.

## Introduction

Plant-pathogenic fungi pose a growing threat to global food security, causing substantial yield losses and driving the emergence of fungicide-resistant populations (1, 2). At the same time, mounting evidence of the environmental and health risks associated with synthetic pesticides is accelerating the transition toward sustainable disease-management approaches, with biological control agents emerging as a leading alternative (3–5).

In this context, the phylum *Actinomycetota*, formerly known as Actinobacteria, has emerged as a promising group for the development of eco-friendly plant disease management strategies, due to their prolific array of bioactive secondary metabolites with antagonistic capacity against phytopathogenic fungi (6–8). Bioprospecting for new biopesticides has focused predominantly on the genus *Streptomyces*, overlooking the vast potential of “rare” actinomycetes. These underexplored taxa have already yielded clinically important antibiotics such as rifamycin (*Amycolatopsis*), erythromycin (*Saccharopolyspora*), and teicoplanin (*Actinoplanes*), and remain an untapped reservoir for novel bioactive compounds (9, 10).

Among *Actinomycetota*, the genus *Saccharomonospora* is a particularly intriguing yet underexplored lineage within the family *Pseudonocardiaceae*, with substantial biosynthetic potential. Recent taxogenomic and comparative genomic analyses showed that *Saccharomonospora* species harbor numerous biosynthetic gene clusters (BGCs) that encode a diverse and novel array of polyketide synthases (PKS) and non-ribosomal peptide synthetases (NRPS) relative to other *Actinomycetota* genera (11). The designation of *Saccharomonospora* and related genera as “rare Actinomycetes” is not due to their scarcity in nature, but their underrepresentation in culture collections, often as a consequence of slow growth rates and the absence of targeted isolation strategies (10, 12).

A recent greenhouse study in our lab revealed that seedlings inoculated with the fungal pathogen *Fusarium oxysporum* harbored a significantly higher relative abundance of *Saccharomonospora* in the rhizosphere of disease-suppressive, organically amended plants than in non-amended controls, suggesting that *Saccharomonospora* may play a role in antagonizing soilborne fungal pathogens (13).

Consequently, this study aimed to investigate the biosynthetic potential and antifungal activity of *Saccharomonospora* species, with the goal of identifying strains capable of producing bioactive compounds that could be harnessed for the biological control of plant pathogens. Initially, we conducted comparative genomic analysis of *Saccharomonospora* isolates and publicly available genomes from NCBI focusing on BGCs predicted to encode compounds with potential antimicrobial activity; and subsequently, evaluated the antifungal capacity of cell-free supernatants from soil-derived *Saccharomonospora* isolates. Three strains significantly inhibited growth of phytopathogenic Fusarium strains, prompting a metabolomics analysis to identify the secreted metabolites potentially linked to the antifungal activity.

## Materials and methods

### Bacteria strains used in this study

The following type strains were purchased from the Leibniz Institute DSMZ (https://www.dsmz.de/): *Saccharomonospora azurea* NA-128 (DSM 44631), *Saccharomonospora caesia* (DSM 43044), *Saccharomonospora cyanea* SIIA 88134 (DSM 44106), *Saccharomonospora glauca* K62 (DSM 43769), and *Saccharomonospora xinjiangensis* strain XJ-54 (DSM 44391). For routine cultivation, these type strains, along with *Saccharomonospora viridis* R81, which was isolated in this study, were grown in tryptic soy broth (TSB) or TSB supplemented with agar for solid media (BD, USA) at 30°C, except for *S. viridis* R81, which was grown at 50°C.

### Isolation, whole-genome sequencing assembly and annotation of Saccharomonospora

*Saccharomonospora* strain R81 was isolated from the rhizosphere soil of compost-amended cucumber plants previously infected with *Fusarium oxysporum f. sp. radicis-cucumerinum* (FORC), as described in our earlier study (13). Isolation was performed on R8 agar medium (14) using dilution streaking, followed by incubation at 50 °C for 5 days. Preliminary taxonomic identification was based on 16S rRNA gene sequencing using primers 11F (GTTTGATCCTGGCTCAG) and 1392R (ACGGGCGGTGTGTRC). PCR products were sequenced by Macrogen Inc. (Seoul, Korea), and the resulting sequences were queried against the NCBI non-redundant (nr) database using BLASTn, with >95% identity used for genus-level assignment.

High-molecular-weight genomic DNA was extracted from cells of strain R81 using the MagAttract HMW DNA Kit (Qiagen, USA) according to the manufacturer’s instructions. Whole-genome sequencing was performed by Plasmidsaurus Inc. (Eugene, OR, USA) using both Oxford Nanopore and Illumina platforms. Hybrid genome assembly was carried out using Unicycler v0.5.0 (15), and genome annotation was conducted using Bakta v1.11.0 (16) with default parameters. The assembled genome was submitted to the Type (Strain) Genome Server (TYGS; https://tygs.dsmz.de) for whole-genome-based taxonomic assignment and species delineation using digital DNA-DNA hybridization (dDDH) (17).

### Retrieval and dereplication and phylogenomic analysis of *Saccharomonospora* species

A total of 35 *Saccharomonospora* genomes from the NCBI RefSeq database were downloaded using a custom script from BacLife (18), and grouped according to source environment reported in the NCBI metadata. To minimize redundancy, we clustered the genomes at a 0.99 Mash-estimated Average Nucleotide Identity (ANI) distance (19) and randomly selected one genome from each cluster to represent the group. The final dataset consisted of 16 genomes from NCBI and *Saccharomonospora viridis* strain R81, which was isolated from a compost-amended soil in this study (**Supplementary Table 1**).

### Analysis of secondary metabolite encoding biosynthetic gene clusters (BGCs)

The selected *Saccharomonospora* genomes were annotated for secondary metabolite encoding BGCs using antiSMASH version 7.1.0 (20) with the flags --cb-general --cb-knownclusters --cb-subclusters --asf --pfam2go --genefinding-tool prodigal --smcog-trees and strictness ‘relaxed’ settings. The identified BGCs were then grouped into gene cluster families (GCFs) using BiG-SCAPE version 1.0.1 (21) with the parameters --include_singletons --mix --no_classify --cutoffs 0.3 0.5 0.7 --clans-off --hybrids-off. The flag --mibig was used to compare BGCs to reference clusters in the MIBiG database. BiG-SCAPE network files were visualized in Cytoscape (Shannon et al., 2003) and used to generate a GCF binary matrix with a custom Python script. A specific BiG-SCAPE analysis for NRPS, PKS, and NRP-PKS hybrid BGCs was performed by selecting all relevant BGCs identified by antiSMASH. Homologous BGCs were visualized using Clinker (23).

### Preparation of bacterial cell-free supernatants (CFS)

Bacterial cell-free supernatants (CFS) were utilized in the *in-vitro* bioassay to evaluate the ability of *Saccharomonospora* strains to secrete extracellular metabolites with antifungal activity. CFS were prepared as follows: Strains were first revived from frozen stocks by streaking onto Tryptic Soy Agar (TSA; Tryptic Soy Broth [TSB], BD Bacto™ Soybean-Casein Digest Medium, supplemented with 1.5% w/v agar) plates, which were incubated for 7 days at 30 °C (50 °C for *S. viridis* R81) to obtain mature single colonies. Starter cultures were then prepared by scraping cells from the 7-day-old TSA plates and inoculating them into 5 mL of TSB, followed by incubation for 3 days at 30 °C (50 °C for *S. viridis* R81) with shaking at 170 rpm. Subsequently, 50 mL of TSB in 250-mL Erlenmeyer flasks were then inoculated with 1 mL of the 3-day-old starter culture, with three flasks per strain. Cultures were incubated at 30°C (50°C for *S. viridis* R81) shaking at 170 rpm, for 7 days. At the end of the incubation period, cultures were centrifuged at 5000 × g for 10 minutes, and the supernatants were filtered through a 0.22 µm Millex-HV syringe filter (Sigma-Aldrich, Israel). The filtered CFS were stored at 4°C for use in the antifungal assays described below.

### Fungal cultures

The GFP-labeled *Fusarium oxysporum f. sp. radicis cucumerinum* (FORC) strain used in this study (24) was kindly provided by the Covo lab at the Hebrew University Faculty of Agriculture, Food and Environment. *Fusarium solani* and *Fusarium oxysporum f. sp. coriandriii* strains were obtained from Dr. Omer Frenkel’s lab at the Agricultural Research Organization, Israel. Fungal cultures were grown on potato dextrose agar (PDA) or potato dextrose broth (PDB, BD, USA). Cultures were stored in 50% glycerol at −80°C and maintained on PDA plates (BD Difco Laboratories, Detroit, MI, USA) supplemented with 250 mg/L chloramphenicol (Thermo Fisher Scientific, Waltham, MA, USA) at 4°C (PDA+ plates). GFP-FORC was initially thawed on PDA supplemented with 100 μg/mL hygromycin (Thermo Fisher Scientific) and then cultured on PDA+ as described in Kraut-Cohen et al. (2024) (25).

### Preparation of fungal spore solution

Spores for *in-vitro* antifungal assays were obtained using a previously documented method (25). Briefly, fungal plugs from PDA+ plates were placed in 15 mL of PDB for FORC cultivation, or in Czapek Dox broth for *F. solani* and *F. coriandrii* cultivation and incubated in an orbital shaker (100 rpm) at 25°C for 5 days. Spores were then extracted by filtering the fungal liquid culture through a nylon cell strainer (mesh size 40 μm; Corning Inc., Corning, NY, USA) and suspended in PDB to achieve a final concentration of 1 × 10⁵ spores/mL.

### Antifungal assays

*In-vitro* antifungal assays were performed for each pathogen by adding 100 µL of CFS to 100 µL of fungal spore suspension (1 × 10⁵ spores/mL) in 96-well plates. CFS-treated fungal spores were incubated at 25°C for 24 hours, and growth was monitored by measuring the optical density (OD) using a multimode microplate reader (Synergy H1, BioTek, USA), which was set to perform an area scan for comprehensive detection across the well surface. Fungal spores amended with 12.5 ppm of the antifungal compound cycloheximide (Sigma-Aldrich) served as a positive control, while spores amended with TSB served as a negative control. Growth was quantified by measuring the change in OD over 24 hours of incubation and normalizing to the initial OD measurements before calculating the area under the growth curve (AUC). Percentage inhibition was calculated as:

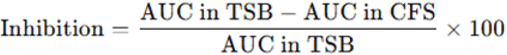

Where **AUC in TSB** refers to the area under the growth curve for fungal spores treated with bacterial TSB medium (negative control), **AUC in CFS** refers to the area under the growth curve for fungal spores treated with the CFS. The result was expressed as a percentage inhibition of fungal growth in the presence of CFS relative to the negative control (TSB).

### Metabolomics

The SPE-cleaned extracts from *S. cyanea*, *S. viridis* R81, *S.xinjiangensis* XJ-54 and TSB media as background were analyzed using Vanquish HPLC system (Thermo Scientific, Waltham, MA, USA) equipped with a Kinetex 1.7 μm C18 reversed phase HPLC column (50 x 2.1 mm) and a *tims*TOFPro2 mass spectrometer (Bruker Daltonics, Bremen, Germany) with an ESI source. Data were acquired in positive and negative ionization mode for parent mass (MS^1^) and tandem MS/MS for molecular fragmentation (MS^2^). For each sample, 1 mg of dried extract was dissolved in methanol, and 1 µL of the resulting solution was injected per LC-MS run. (4-(3,5-dimethyl-1H-pyrazol-4-yl)methyl)-5-methylisoxazol-3-yl)(4-(pyrazolo[1,5-a]pyrimidin-7-yl)piperidin-1-yl)methanone (PharmBricks, Israel) was used as an internal standard (retention time: 10.0 min). The following gradient was used for chromatographic separation: start from 1% of B (Acetonitrile + 0.1% Formic acid) and 99% of A (H_2_O + 0.1% Formic acid) to 5% B in 2 min, 5 to 10% B for 3 min, 10 to 40% B for 4 min, 40–95% B in 1 min, held at 95% B for 0.5 min, 95–1% B in 0.5 min and kept at 1% B for 4 min at a flow rate of 0.3 mL/min throughout the run. The MS analysis was performed on *tims*TOFPro2 mass spectrometer (Bruker Daltonics, Bremen, Germany) equipped with ESI source.

The MetaboScape 2023b software was used for untargeted metabolite profiling, with auto MS/MS matching performed on the generated bucket table. Calibration of the MS data was conducted using sodium formate, and the analysis was carried out according to the parameters provided in **Supplementary Table 2**. Annotation of bucket table was produced based on spectral libraries (uploaded from MassBank of North America (MoNA)) and target lists (SuCComBase-NPAtlas-Food DB and HMDB (obtained from the Bruker company)). The analyte list matches were determined based on the following criteria: m/z ± 2/50 ppm, retention time ± 0.1/0.5 minutes, MS/MS score ≥ 900/600, mSigma ≥ 20/100, and CCS ≥ 1/3.

A molecular network of the all the identified metabolites in all samples was performed with the web-based GNPS pipeline (26) using previously validated parameters and chemical class prediction was performed using CANOPUS (27).

### Statistical analysis

Statistical comparison of gene cluster family (GCF) abundance was performed using one-way ANOVA, with multiple comparisons of means conducted using the Tukey–Kramer (HSD) test (*p* < 0.05). Statistical differences in GCF composition, based on the GCF binary matrix from BiG-SCAPE, were assessed using PERMANOVA with Jaccard distance, 999 permutations, and Benjamini-Hochberg (BH) correction of p-values, using Vegan package in R. Statistical comparisons of fungal inhibition by different *Saccharomonospora* strains were performed using the Kruskal-Wallis test (*p* < 0.05), followed by post-hoc pairwise comparisons in JMP software (JMP®, Version 17. SAS Institute Inc., Cary, NC, 1989-2023).

## Results

### Isolation and *in-silico* phylogenetic assignment of *Saccharomonospora* isolate R81

Recent computational analyses showed that *Saccharomonospora* possesses novel and diverse biosynthetic gene clusters (11), and our previous work demonstrated that this genus was significantly enriched in rhizospheres where *Fusarium oxysporum* disease suppression was observed under organic amendment (13). These findings motivated the targeted isolation of *Saccharomonospora* from these rhizosphere samples for genomic and phenotypic characterization.

One isolate forming blue-green colonies was recovered on R8 agar and identified as *Saccharomonospora* based on full-length 16S rRNA gene sequencing and whole-genome analysis. Digital DNA-DNA hybridization (dDDH) analysis via the Type Strain Genome Server (TYGS) revealed that strain *Saccharomonospora* R81 exhibited high genomic similarity to the type strain *Saccharomonospora viridis* DSM 43017, with dDDH values of 97.5% (d0), 96.1% (d4), and 98.4% (d6). Similar values were obtained when compared to S. *viridis* ATCC 33517 (96.6%, 96.0%, and 97.9%, respectively). These values are well above the 70% threshold for species delineation, and the G+C content differences were minimal (0.1-0.15%), confirming that strain R81 belongs to the species *S. viridis*. The isolate is hereafter referred to as *S. viridis* R81 (NCBI accession: PRJNA1307827).

### *Saccharomonospora* CFS exhibit broad variation in antifungal efficacy against *Fusarium* phytopathogens

We evaluated the in-vitro antifungal activity of cell-free supernatants (CFS) from *Saccharomonospora viridis* R81, soil-derived type strains *S. azurea*, *S. cyanea*, *S. caesia*, *S. xinjiangensis*, and the compost-derived type strain *S. glauca* against three Fusarium phytopathogens: *Fusarium oxysporum* FORC, *F. Solani* and *F. oxysporum f. sp. coriandriii* (FCOR), using cycloheximide (12.5 ppm) as a positive control for fungal inhibition and bacteria-free tryptic soy broth (TSB) as a negative control (**Figure 1**).

**Figure 1.**
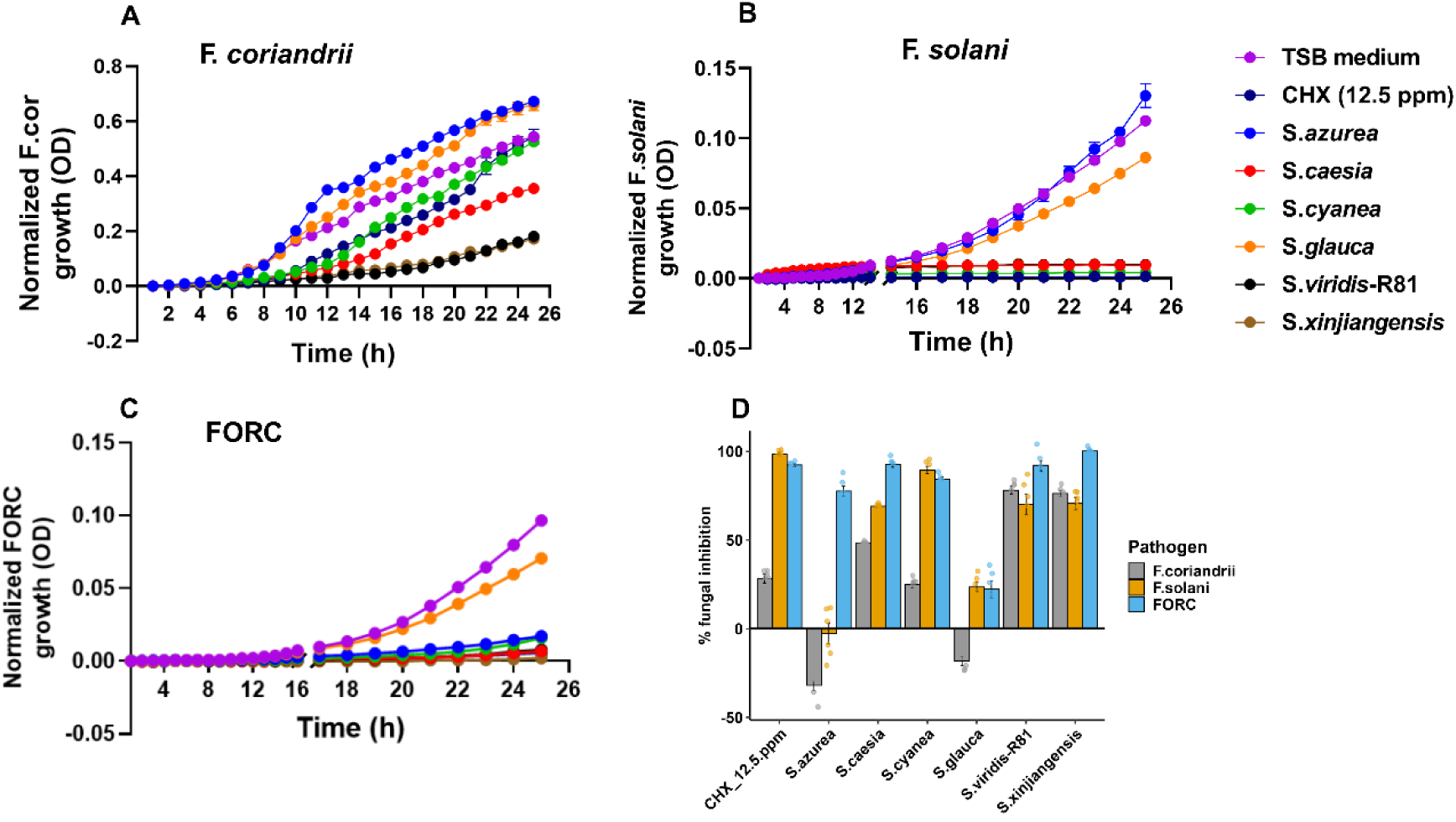
*S. xinjiangensis* and *S. viridis* R81 CFS exhibit broad-spectrum antifungal activity against Fusarium phytopathogens. Growth kinetics of *F. coriandrii* **(A)**, *F. solani* **(B)** and FORC **(C)** over 24 h when treated with cell-free supernatants (CFS) from six *Saccharomonospora* strains compared to TSB medium (negative control) and cycloheximide (CHX, 12.5 ppm; positive control). Growth was quantified by the change in Optical density at 600 nm (OD₆₀₀) monitored over 24h of incubation and normalized relative to the initial OD measurements before estimation of the area under the growth curve (AUC), and percentage of inhibition was calculated based on AUC. **(D)** Bar plots showing percentage of inhibition of *F. coriandrii* (gray bars), *F.solani* (orange bars) and FORC (blue bars) relative to the negative control. Data represent the mean of six technical replicates. In Panels A-C, curves show mean fungal growth, and error bars indicate the standard error of the mean (SEM). In Panel D, bars represent mean percent inhibition, error bars show SEM, and individual dots correspond to the six replicate measurements.

All *Saccharomonospora* strains exhibited antifungal activity, although the degree of inhibition varied across the three Fusarium pathogens. All strains antagonized FORC, with values ranging from 22% inhibition in *S. glauca* to 100% in *S. xinjiangensis*. Five of the six strains antagonized *F. solani*, with inhibition ranging from 24% to 90%. *S. cyanea* showed the highest activity, whereas S. *azurea* exhibited no inhibition. The pathogen *F. coriandrii* was the least susceptible, with two strains showing weak or no inhibition (*S. azurea*, -32%; *S. glauca*, -18%). However, *S. xinjiangensis* (∼76%) and *S. viridis* R81 (∼78%) displayed strong inhibitory activity, exceeding CHX (only ∼28% inhibition). Collectively, *S. xinjiangensis* and *S. viridis* R81 produced the most potent and broad-spectrum CFS, capable of effectively antagonizing multiple Fusarium species and outperforming the commercial antifungal compound in certain cases.

### Strain-specific BGC repertoires likely underpin variation in antifungal activity

To examine the genomic basis of the observed antifungal activity, we analyzed the secondary metabolite biosynthetic potential of the five strains with available genome sequences (*S. caesia* was excluded due to the absence of a genome in NCBI).

Using antiSMASH, we identified 58 BGCs across the five strains, with *S. viridis* R81 harboring the least (9 BGCs), and *S. cyanea* and *S. xinjiangensis* containing the highest count with 14 BGCs. BiG-SCAPE software was used to compare these identified BGCs against previously characterized clusters archived in the MIBiG database, and a sequence similarity network of the identified BGCs was constructed to visualize the biosynthetic diversity within our isolates. A small core set of BGCs encoding for ectoine, arylpolyene, and terpene was present in all strains irrespective of their antifungal activity (**Figure 2**), indicating that these conserved clusters are unlikely to explain variations in their antifungal activity. In contrast, 19 BGCs were strain-specific and showed no similarity to any MiBIG reference BGC, highlighting that these strains harbor novel biosynthetic pathways (**Figure 2**). *S. viridis* R81, which exhibited strong and broad antifungal activity, uniquely encoded taromycin- and clavulinic acid-like BGCs in addition to a mirubactin-like siderophore cluster, representing plausible genetic sources of bioactive metabolites. *S. xinjiangensis*, which similarly showed strong antifungal activity, contained 14 BGCs, none of which matched closely to MiBIG reference clusters, suggesting that its bioactivity arises from uncharacterized biosynthetic pathways.

**Figure 2.**
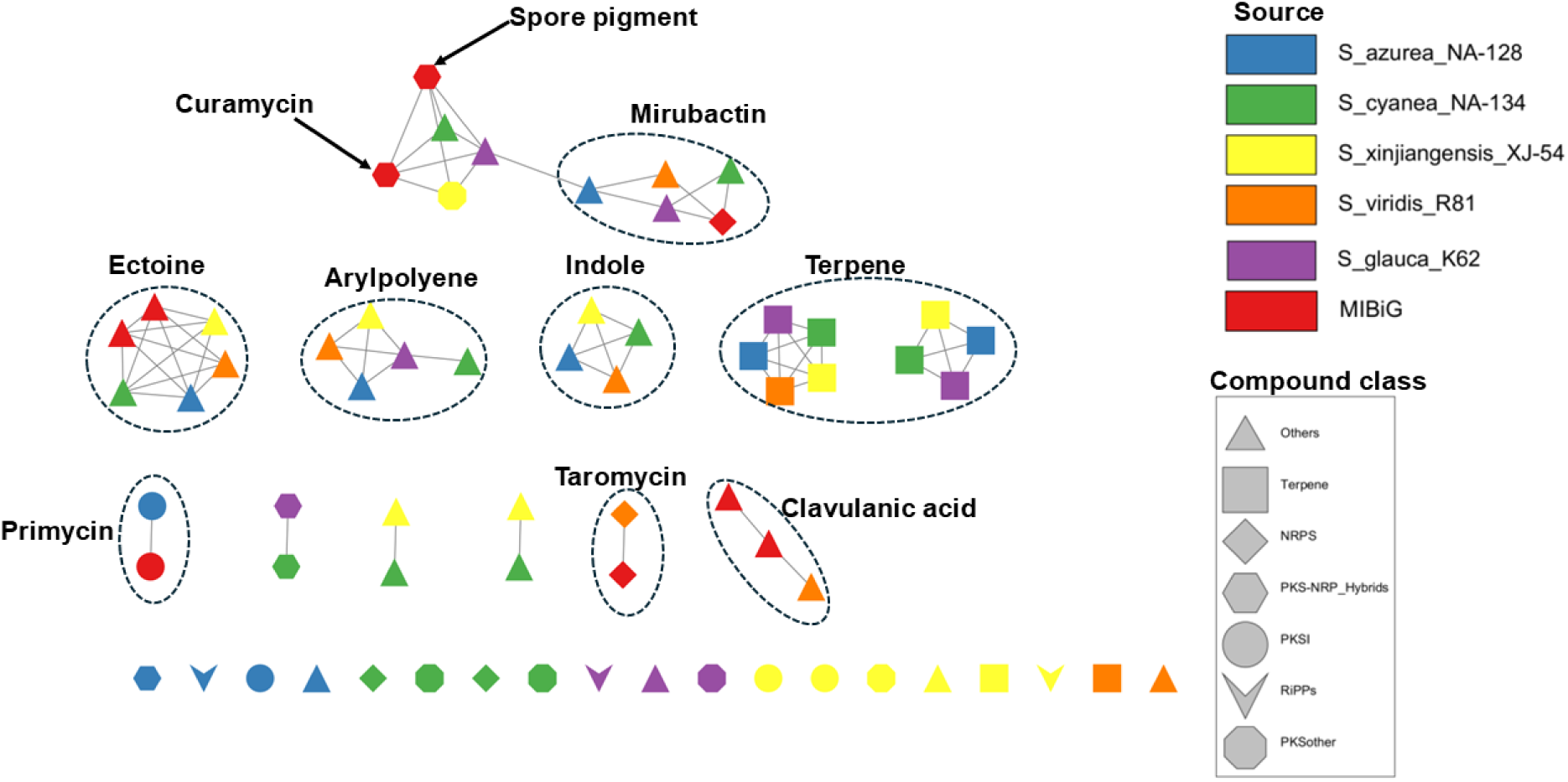
Network analysis of antifungal-tested *Saccharomonospora* strains revealed largely uncharacterized biosynthetic potential. The BGC network was constructed using BiG-SCAPE with a 0.5 similarity cutoff and includes BGCs from the five indicated strains. Reference BGCs from the MIBiG database are shown as red nodes. Nodes connected to these reference nodes indicate BGCs with sequence similarity to characterized pathways and likely encode related compound classes. Node shape denotes BGC class, and node color denotes the strain of origin.

Surprisingly, *S. azurea* which harbored a type I polyketide synthase (T1PKS) BGC homologous to a gene cluster encoding a macrolide lactone antibiotic primycin, exhibited weak to no antifungal activity, highlighting that BGC presence does not necessarily translate to expression or production of the encoded compound at titers sufficient for bioactivity under the tested conditions.

Collectively, these patterns indicate that variation in antifungal activity among these strains likely reflects differences in their strain-specific BGC repertoires, with both known antibiotic-associated pathways and uncharacterized clusters representing plausible genetic determinants of the observed phenotypes.

### Comparative metabolomics reveals expanded biosynthetic repertoires and enhanced metabolite production in *S. viridis* R81 and *S. xinjiangensis* compared to *S. cyanea*

Genetic potential alone does not establish which metabolites are produced, and therefore we characterized the metabolomes of the two most active strains (*S. xinjiangensis* and *S. viridis* R81) and a moderately active strain (*S. cyanea*) using untargeted LC-MS/MS. Unsupervised PCA clustering and pairwise PERMANOVA did not reveal significant differences in metabolite composition among the three strains (**Supplementary Figure 1**). However, metabolite overlap analysis highlighted shared and strain-specific features (**Figure 3A**). *S. viridis* R81 and *S. xinjiangensis*, which exhibited broader antifungal activity, shared a greater number of metabolites (25%, 58/233) and possessed more unique features (34 and 25, respectively) than *S. cyanea* (5), despite *S. xinjiangensis* being more closely related to *S. cyanea* than to *S. viridis* R81. Although 40% (93/233) of metabolites were shared among the three strains, the analysis of metabolite intensities revealed substantial quantitative differences within this common metabolome. *S. viridis* R81 and *S. xinjiangensis* exhibited higher overall intensities across most metabolite classes than *S. cyanea*. Of these shared metabolites, only 48 (∼52%) were successfully assigned to chemical classes using CANOPUS annotation (27). Within the shared metabolome, classes such as ornithine and nicotinic acid alkaloids, linear polyketides, coumarins, fatty esters, and fatty amides were more abundant in *S. viridis* R81 and *S. xinjiangensis* (**Figure 3B**), and it is possible that these quantitative differences are likely associated with enhanced antifungal activity of these two strains.

**Figure 3.**
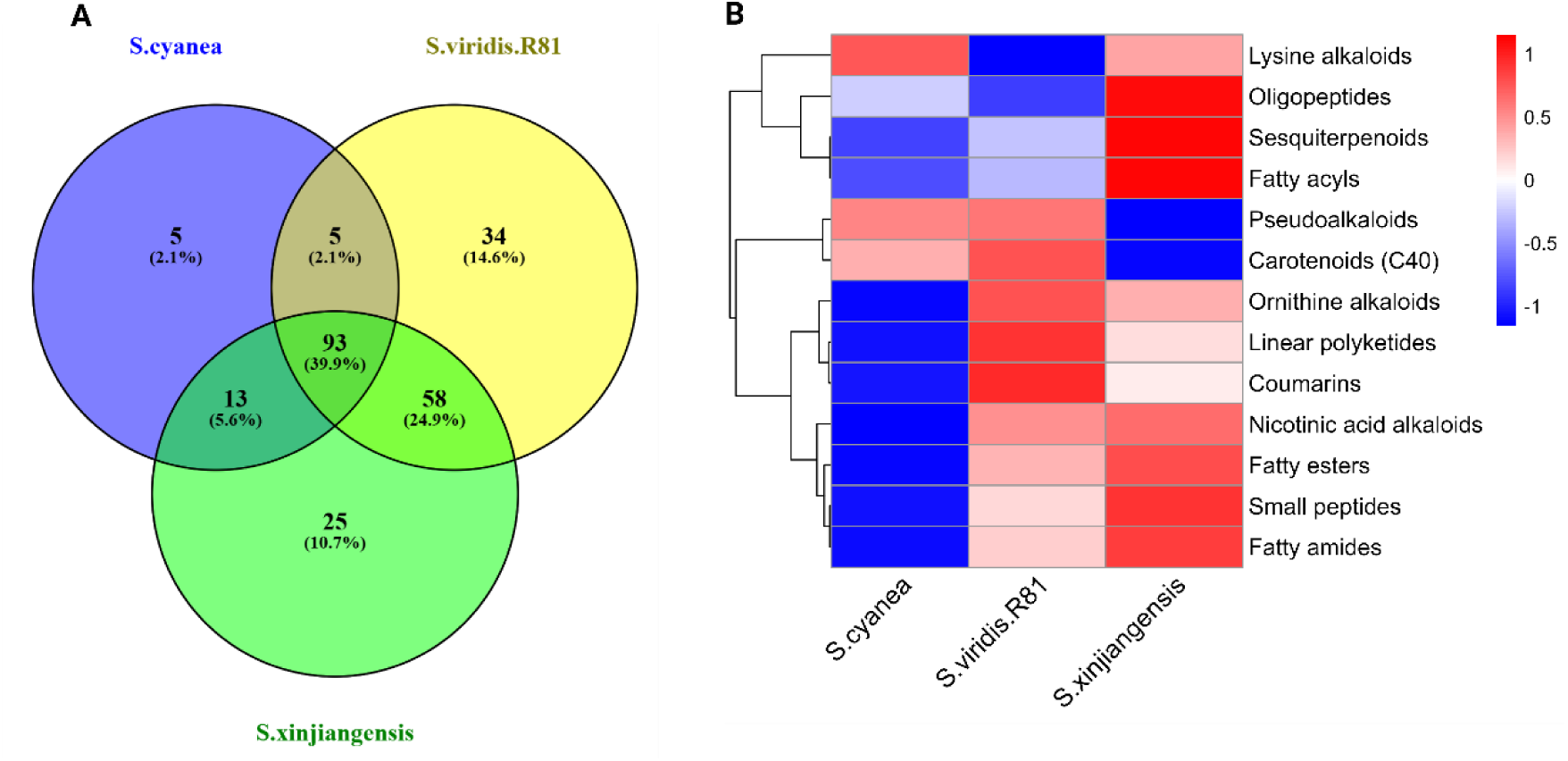
*S. viridis* R81 and *S. xinjiangensis* display richer metabolite profiles and show higher abundances of the shared metabolites compared to *S. cyanea*. **(A)** Venn diagram showing unique and shared metabolite features in *S. cyanea*, *S. viridis R81*, and *S. xinjiangensis*. **(B)** Heatmap showing relative abundance of 13 metabolite classes within the shared metabolome that was classified by CANOPUS. Values represent the mean abundance of features within each class, averaged across three replicates per strain log₁₀-transformed, and shown as row-wise Z-scores. Red indicates higher and blue lower abundance.

Of the 58 metabolites shared between *S. viridis* R81 and *S. xinjiangensis*, 25 could be annotated, and these spanned diverse chemical classes (**Supplementary Table 3**). The majority were alkaloids and peptide derivatives, including pyridine, quinolizidine, piperidine, and purine alkaloids, as well as several dipeptides and tripeptides. Additional groups that were enriched included polyamines, fatty esters, fatty alcohols, open-chain polyketides, coumarins, and terpenoid derivatives. Most of these metabolites showed markedly higher intensities in one or both of the two strains and were absent in *S. cyanea*. Alkaloid-derived metabolites are frequently linked to antifungal bioactivity, and thus their enrichment in the CFS of the two most potent *Saccharomonospora* strains is highly noteworthy in the context of the observed Fusarium suppression.

Collectively, these findings suggest that the broad antifungal activity of *S. viridis*-R81 and *S. xinjiangensis* may not only result from the production of unique metabolites, but also from elevated production of shared metabolites, particularly alkaloids, polyketides, and peptide derivatives.

### BGC composition, not abundance, varies among *Saccharomonospora* strains isolated from different environments

After identifying strain-specific BGCs linked to antifungal activity in our focal isolates, we surveyed all publicly available *Saccharomonospora* genomes to characterize the genus-wide biosynthetic landscape. We compiled 16 non-redundant genomes spanning diverse reported isolation environments (**Supplementary Table 1**). Fourteen of these were assigned to three broad environmental categories, soil (six strains, including *S. viridis* R81), hypersaline soil (four strains), and aquatic habitats (four strains from marine, lake, or fishpond sediments), while two additional genomes from compost and plant seed were included in the dataset but not in group-level comparisons due to limited representation.

BGC annotation using antiSMASH predicted 177 BGCs across the 14 representative genomes, which clustered into 89 GCFs. GCF abundance, estimated as the number of GCFs normalized to genome size for each genome did not significantly differ between the three environments (one-way ANOVA, Tukey’s post-hoc test, *P* > 0.05) (**Figure 4A**). In contrast, ordination based on Jaccard distances of GCF presence/absence revealed distinct and significant differences between soil-derived genomes, and those from the aquatic and hypersaline saline environments (adjusted *P* = 0.012, R² = 0.24) (**Figure 4B**).

**Figure 4.**
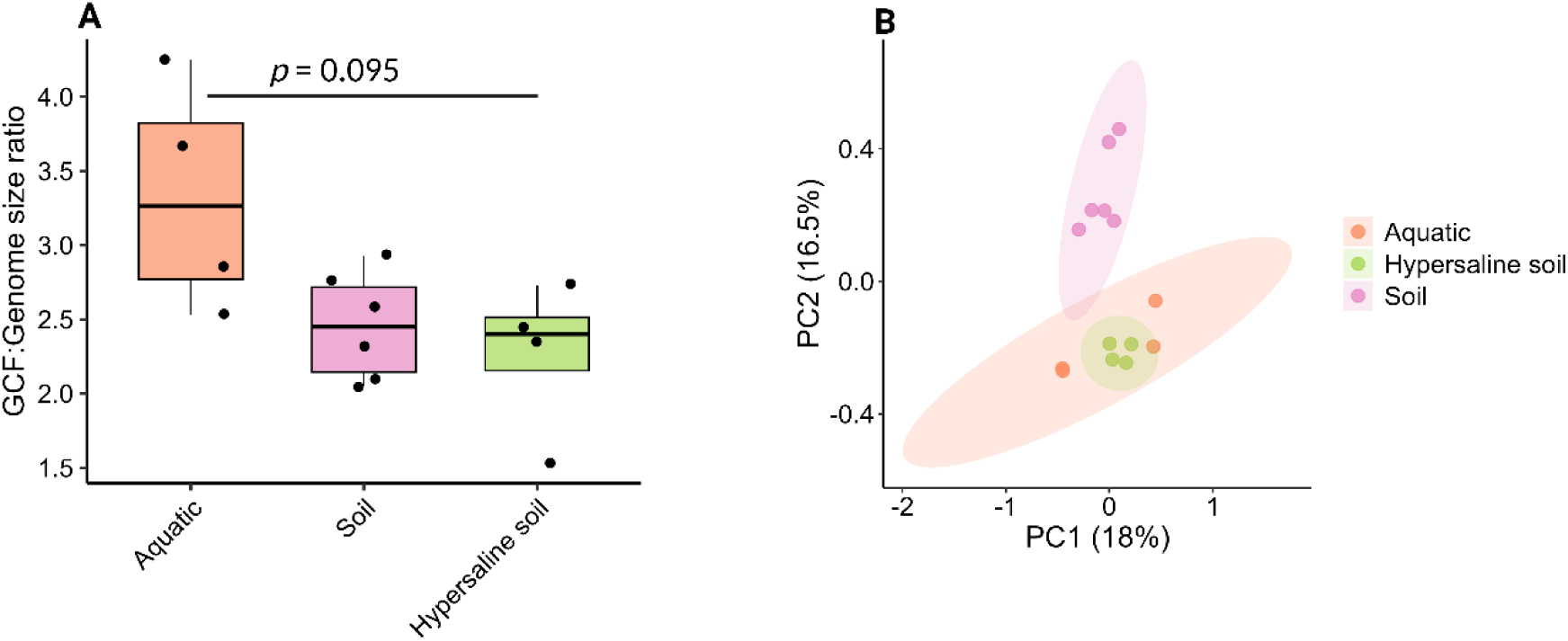
*Saccharomonospora* strains differ in BGC composition but not in overall BGC abundance. (A) GCF richness normalized to genome size across *Saccharomonospora* strains from different isolation sources. No statistical significance was observed between environments (Kruskal-Wallis *P* = 0.095). (B) PCA analysis of *Saccharomonospora* GCFs in the three evaluated environments.

### *Saccharomonospora* possess diverse, novel BGCs with lineage-associated conservation

The 117 BGCs identified in the 14 analyzed *Saccharomonospora* genomes were classified into 11 distinct classes, with terpene (32) and RiPPs (25) being the most abundant class (**Figure 5A**). The “Others” category, comprising 37 BGCs, included clusters that could not be confidently assigned to any known BGC class by antiSMASH or BiG-SCAPE. BGC clustering of all the 117 BGCs into gene cluster families (GCFs) revealed that the majority of GCFs were unique to individual strains. Specifically, 47 GCFs were singletons (present in only one strain), while only 3 were shared by 9-14 strains, reflecting the high biosynthetic diversity and low level of GCF conservation within the genus (**Figure 5B**).

**Figure 5.**
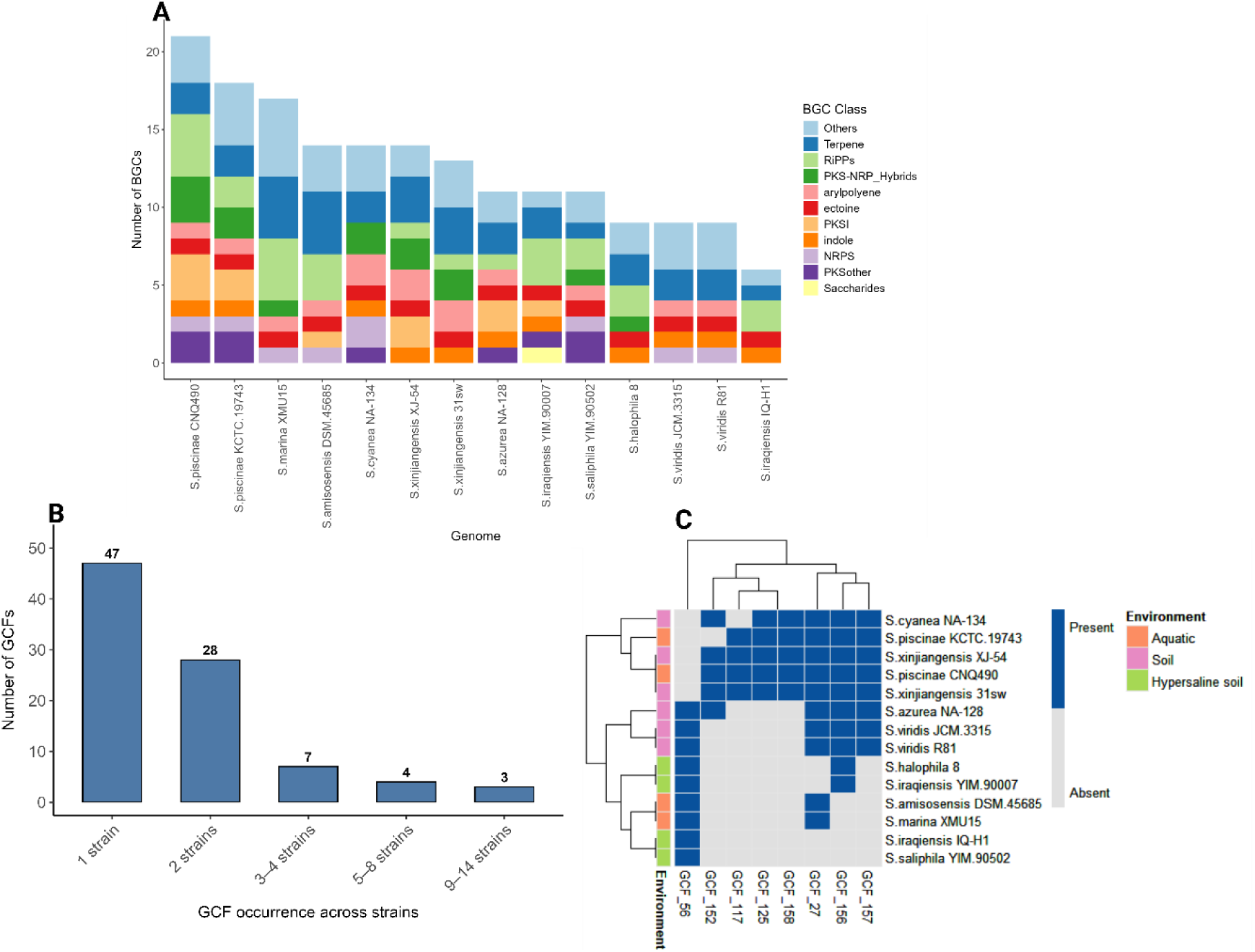
*Saccharomonospora* strains encode diverse, strain-specific BGC repertoires. (**A**) BGC class composition per genome as predicted by antiSMASH. Genomes were ordered by total BGC count. (**B**) Distribution of GCFs based on their occurrence across *Saccharomonospora* strains. Most GCFs are singletons, occurring in only one strain. (**C**) Presence-absence matrix of GCFs found in more than three strains, showing how they are shared across *Saccharomonospora* strains isolated from aquatic, soil, and hypersaline environments. Clustering is based on GCF presence.

We further investigated whether biosynthetic gene cluster families in *Saccharomonospora* strains are conserved according to environmental origin. Analyzing only GCFs present in more than three genomes revealed phylogenetic rather than environment-associated clustering (**Figure 5C**), indicating that biosynthetic repertoires are not shaped by environment. Although three GCFs (27, 156, 157) appeared in all soil strains, their presence in two aquatic *S. piscinae* strains suggests phylogenetic retention rather than environment.

To assess the novelty of BGCs, a biosynthetic novelty index (NI) was calculated. For each GCF, the novelty index was defined as:

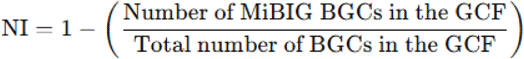

NI reflects the proportion of uncharacterized BGCs within each GCF, where higher values indicate greater biosynthetic novelty. For instance, GCF_2504, which included two BGCs, one corresponding to the primycin BGC from the MiBIG database and one from *S. azurea*, had a novelty index (NI) of 0.5, indicating that half of the BGCs in this GCF corresponded to previously characterized biosynthetic pathways. Genome-level novelty scores were obtained by averaging the NI values across all GCFs contributed by each genome.

All strains regardless of the isolation environment, exhibited high mean novelty scores, ranging from 0.828 to 0.977 (**Supplementary Table 4**), indicating that the majority of their BGCs belong to novel or poorly characterized GCFs. These results underscore the untapped biosynthetic potential of *Saccharomonospora*, with most strains harboring BGCs belonging to GCFs with limited representation in current natural product databases.

## Discussion

Within the phylum *Actinomycetota*, the genus *Streptomyces* remains the best-characterized source of bioactive secondary metabolites, contributing many clinically and agriculturally relevant natural products, including antibiotics and antifungals (9, 10). However, a growing body of evidence indicates that less represented “rare” actinomycete genera, such as *Saccharomonospora*, *Micromonospora*, and *Nocardiopsis*, also harbor extensive and largely uncharacterized biosynthetic potential (11, 28). Here, we explored the biosynthetic capacity and antifungal potential of *Saccharomonospora* species, testing the hypothesis that soil-derived strains produce metabolites that antagonize soilborne fungal pathogens as a result of their co-occurrence with soilborne phytopathogenic fungi in soil environments.

Nearly all of the screened CFS inhibited at least one of the three tested Fusarium pathogens, demonstrating the potential of *Saccharomonospora* as a source of potent antifungal compounds. The *S. xinjiangensis* and *S. viridis* R81 strains exhibited the broadest antifungal spectrum and strongest inhibition, effectively suppressing all three Fusarium strains and in some cases outperforming cycloheximide. The observed antifungal activity likely arises from the production of bioactive secondary metabolites, some encoded by BGCs, that constitute the chemical basis of antagonism in actinomycetes (29–31).

While BGC composition in soil-derived genomes showed distinct separation from aquatic and hypersaline environments, we found no strong evidence that environment determines BGC repertoire. No GCF was exclusive to any habitat, and most were strain-specific. The limited number of available genomes per environment may have constrained detection of rare environment-associated BGC signatures.

Interestingly, most BGCs especially NRPS and PKS were unique to individual strains reflecting potential for high chemical diversity. These BGCs often encode bioactive metabolites, such as antibiotics, siderophores, or toxins, (32, 33) whose ecological benefits are context-dependent, targeting specific competitors or hosts and might be important for adaptation to specific environments rather than survival (34, 35).

No major differences were observed in the GC content of BGC relative to whole-genomes, supporting vertical inheritance rather than recent horizontal gene transfer events. This is consistent with observations that vertical descent and diversification can drive species-level BGC variation (36). Although we did not examine BGC evolutionary trajectories in detail, our results suggest that lineage-specific retention and divergence probably underlie the observed BGC distribution in *Saccharomonospora*. Similar strain-specific BGC diversification has been observed in other actinomycetes and symbiotic bacteria (36–38), indicating that such lineage-specific patterns are a common feature of specialized metabolite evolution.

Conversely, indole, ectoine, arylpolyene, and terpene BGCs were nearly ubiquitous across *Saccharomonospora* genomes. The widespread conservation of these pathways suggests that they encode either ancient biosynthetic functions retained across the lineage and/or core traits essential for survival. These pathways are well known for their roles in stress tolerance, oxidative protection, and ecological signaling (39–41), essential for survival in the diverse environments inhabited by *Saccharomonospora*.

The *Saccharomonospora* genomes analyzed in this study provide strong evidence of highly novel biosynthetic potential. To date, only two compounds from this genus have been experimentally characterized: the macrolide antibiotic primycin from *S. azurea* (42), and the cyclic lipopeptide taromycin from *S. viridis* CNQ-490 (43). The predominance of uncharacterized, strain-specific BGCs underscores *Saccharomonospora* as an untapped source of chemically diverse metabolites with potential for biotechnological exploitation, including the development of biopesticides to combat soilborne plant pathogens (35).

The shared metabolome of the three most inhibitory *Saccharomonospora* strains was enriched in polyketides, coumarins, and alkaloids, which may collectively underpin the antifungal activity observed. This pattern is consistent with the genomic detection of at least one polyketide synthase (PKS) or hybrid PKS-NRPS biosynthetic gene cluster in every strain, and the presence of a conserved indole BGC across all genomes. Antifungal activity of coumarins and alkaloids against a wide range of plant pathogenic fungi has been previously reported (44, 45). In particular, Sequin et al. (2023) reported that the alkaloid fraction from *Neltuma nigra* leaves strongly inhibited agriculturally important pathogens, *Cercospora kikuchii*, *C. sojina*, and *Sclerotium rolfsii*, exhibiting lower toxicity than an azole fungicide (46). These findings underscore the promise of natural compounds as effective and environmentally safer alternatives for managing fungal diseases in crops.

In summary, this study expands the understanding of secondary metabolism in “rare” Actinobacteria, revealing *Saccharomonospora* as a source of diverse and unique NRPS, PKS, and hybrid pathways, with several soil-derived strains showing strong *in-vitro* suppression of Fusarium pathogens. These findings highlight the potential of this genus as a robust reservoir for antifungal agents, offering a strategic pathway for sustainable plant disease management. Specifically, the genomic and metabolomic data presented here provide a foundation for discovering novel metabolites with activity against *Fusarium* and other phytopathogens. Isolating and characterizing these bioactive compounds, combined with targeted BGC manipulation to establish gene-metabolite links, will facilitate the discovery of new natural products and advance biocontrol strategies for sustainable plant disease management.

## References

1. Yin Y, Miao J, Shao W, Liu X, Zhao Y, Ma Z. 2023. Fungicide Resistance: Progress in Understanding Mechanism, Monitoring, and Management. Phytopathology 113:707–718.

2. Stukenbrock E, Gurr S. 2023. Address the growing urgency of fungal disease in crops. Nature 617:31–34.

3. Sarrocco S. 2023. Biological Disease Control by Beneficial (Micro)Organisms: Selected Breakthroughs in the Past 50 Years. Phytopathology 113:732–740.

4. Schneider K, Barreiro-Hurle J, Rodriguez-Cerezo E. 2023. Pesticide reduction amidst food and feed security concerns in Europe. Nat Food 4:746–750.

5. Wan NF, Fu L, Dainese M, Kiær LP, Hu YQ, Xin F, Goulson D, Woodcock BA, Vanbergen AJ, Spurgeon DJ, Shen S, Scherber C. 2025. Pesticides have negative effects on non-target organisms. Nat Commun 16:1–16.

6. Palaniyandi SA, Yang SH, Zhang L, Suh JW. 2013. Effects of actinobacteria on plant disease suppression and growth promotion. Appl Microbiol Biotechnol 97:9621–9636.

7. Boubekri K, Soumare A, Mardad I, Lyamlouli K, Ouhdouch Y, Hafidi M, Kouisni L. 2022. Multifunctional role of Actinobacteria in agricultural production sustainability: A review. Microbiol Res 261:127059.

8. Kaur T, Khanna K, Sharma S, Manhas RK. 2023. Mechanistic insights into the role of actinobacteria as potential biocontrol candidates against fungal phytopathogens. J Basic Microbiol 63:1196–1218.

9. Ding T, Yang LJ, Zhang WD, Shen YH. 2019. The secondary metabolites of rare actinomycetes: Chemistry and bioactivity. RSC Adv 9:21964–21988.

10. Parra J, Beaton A, Seipke RF, Wilkinson B, Hutchings MI, Duncan KR. 2023. Antibiotics from rare actinomycetes, beyond the genus Streptomyces. Curr Opin Microbiol 76:102385.

11. Ramírez-Durán N, de la Haba RR, Vera-Gargallo B, Sánchez-Porro C, Alonso-Carmona S, Sandoval-Trujillo H, Ventosa A. 2021. Taxogenomic and Comparative Genomic Analysis of the Genus Saccharomonospora Focused on the Identification of Biosynthetic Clusters PKS and NRPS. Front Microbiol 12.

12. Ezeobiora CE, Igbokwe NH, Amin DH, Enwuru N V., Okpalanwa CF, Mendie UE. 2022. Uncovering the biodiversity and biosynthetic potentials of rare actinomycetes. Futur J Pharm Sci 8.

13. Amutuhaire H, Faigenboim-Doron A, Kraut-Cohen J, Friedman J, Cytryn E. 2025. Identifying rhizosphere bacteria and potential mechanisms linked to compost suppressiveness towards Fusarium oxysporum. Environ Microbiome 20.

14. Amner W, Edwards C, McCarthy AJ. 1989. Improved medium for recovery and enumeration of the farmer’s lung organism, Saccharomonospora viridis. Appl Environ Microbiol 55:2669–2674.

15. Wick RR, Judd LM, Gorrie CL, Holt KE. 2017. Unicycler: Resolving bacterial genome assemblies from short and long sequencing reads. PLoS Comput Biol 13:1–22.

16. Schwengers O, Jelonek L, Dieckmann MA, Beyvers S, Blom J, Goesmann A. 2021. Bakta: Rapid and standardized annotation of bacterial genomes via alignment-free sequence identification. Microb Genomics 7.

17. Meier-Kolthoff JP, Auch AF, Klenk HP, Göker M. 2013. Genome sequence-based species delimitation with confidence intervals and improved distance functions. BMC Bioinformatics 14.

18. Guerrero-Egido G, Pintado A, Bretscher KM, Arias-Giraldo LM, Paulson JN, Spaink HP, Claessen D, Ramos C, Cazorla FM, Medema MH, Raaijmakers JM, Carrión VJ. 2024. bacLIFE: a user-friendly computational workflow for genome analysis and prediction of lifestyle-associated genes in bacteria. Nat Commun 15.

19. Ondov BD, Treangen TJ, Melsted P, Mallonee AB, Bergman NH, Koren S, Phillippy AM. 2016. Mash: Fast genome and metagenome distance estimation using MinHash. Genome Biol 17:1–14.

20. Blin K, Shaw S, Augustijn HE, Reitz ZL, Biermann F, Alanjary M, Fetter A, Terlouw BR, Metcalf WW, Helfrich EJN, Van Wezel GP, Medema MH, Weber T. 2023. AntiSMASH 7.0: New and improved predictions for detection, regulation, chemical structures and visualisation. Nucleic Acids Res 51:W46–W50.

21. Navarro-Muñoz JC, Selem-Mojica N, Mullowney MW, Kautsar SA, Tryon JH, Parkinson EI, De Los Santos ELC, Yeong M, Cruz-Morales P, Abubucker S, Roeters A, Lokhorst W, Fernandez-Guerra A, Cappelini LTD, Goering AW, Thomson RJ, Metcalf WW, Kelleher NL, Barona-Gomez F, Medema MH. 2020. A computational framework to explore large-scale biosynthetic diversity. Nat Chem Biol 16:60–68.

22. Shannon, P., Markiel, A., Ozier, O., Baliga, N. S., Wang, J. T., Ramage, D., Amin, N., Schwikowski, B., and Ideker, T. (2003). Cytoscape: A Software Environment for Integrated Models. Genome Research, 13(11), 2498–2504. 10.1101/gr.1239303.

23. van den Belt M, Gilchrist C, Booth TJ, Chooi YH, Medema MH, Alanjary M. 2023. CAGECAT: The CompArative GEne Cluster Analysis Toolbox for rapid search and visualisation of homologous gene clusters. BMC Bioinformatics 24:1–8.

24. Cohen R, Orgil G, Burger Y, Saar U, Elkabetz M, Tadmor Y, Edelstein M, Belausov E, Maymon M, Freeman S, Yarden O. 2015. Differences in the responses of melon accessions to fusarium root and stem rot and their colonization by Fusarium oxysporum f. sp. radicis-cucumerinum. Plant Pathol 64:655–663.

25. Kraut-Cohen J, Frenkel O, Covo S, Marcos-Hadad E, Carmeli S, Belausov E, Minz D, Cytryn E. 2024. A pipeline for rapidly evaluating activity and inferring mechanisms of action of prospective antifungal compounds. Pest Manag Sci 80:2804–2816.

26. Wang M, Carver JJ, Phelan V V., Sanchez LM, Garg N, Peng Y, Nguyen DD, Watrous J, Kapono CA, Luzzatto-Knaan T, Porto C, Bouslimani A, Melnik A V., Meehan MJ, Liu WT, Crüsemann M, Boudreau PD, Esquenazi E, Sandoval-Calderón M, Kersten RD, Pace LA, Quinn RA, Duncan KR, Hsu CC, Floros DJ, Gavilan RG, Kleigrewe K, Northen T, Dutton RJ, Parrot D, Carlson EE, Aigle B, Michelsen CF, Jelsbak L, Sohlenkamp C, Pevzner P, Edlund A, McLean J, Piel J, Murphy BT, Gerwick L, Liaw CC, Yang YL, Humpf HU, Maansson M, Keyzers RA, Sims AC, Johnson AR, Sidebottom AM, Sedio BE, Klitgaard A, Larson CB, Boya CAP, Torres-Mendoza D, Gonzalez DJ, Silva DB, Marques LM, Demarque DP, Pociute E, O’Neill EC, Briand E, Helfrich EJN, Granatosky EA, Glukhov E, Ryffel F, Houson H, Mohimani H, Kharbush JJ, Zeng Y, Vorholt JA, Kurita KL, Charusanti P, McPhail KL, Nielsen KF, Vuong L, Elfeki M, Traxler MF, Engene N, Koyama N, Vining OB, Baric R, Silva RR, Mascuch SJ, Tomasi S, Jenkins S, Macherla V, Hoffman T, Agarwal V, Williams PG, Dai J, Neupane R, Gurr J, Rodríguez AMC, Lamsa A, Zhang C, Dorrestein K, Duggan BM, Almaliti J, Allard PM, Phapale P, Nothias LF, Alexandrov T, Litaudon M, Wolfender JL, Kyle JE, Metz TO, Peryea T, Nguyen DT, VanLeer D, Shinn P, Jadhav A, Müller R, Waters KM, Shi W, Liu X, Zhang L, Knight R, Jensen PR, Palsson B, Pogliano K, Linington RG, Gutiérrez M, Lopes NP, Gerwick WH, Moore BS, Dorrestein PC, Bandeira N. 2016. Sharing and community curation of mass spectrometry data with Global Natural Products Social Molecular Networking. Nat Biotechnol 34:828–837.

27. Dührkop K, Nothias LF, Fleischauer M, Reher R, Ludwig M, Hoffmann MA, Petras D, Gerwick WH, Rousu J, Dorrestein PC, Böcker S. 2021. Systematic classification of unknown metabolites using high-resolution fragmentation mass spectra. Nat Biotechnol 39:462–471.

28. González-Salazar LA, Quezada M, Rodríguez-Orduña L, Ramos-Aboites H, Capon RJ, Souza-Saldívar V, Barona-Gomez F, Licona-Cassani C. 2023. Biosynthetic novelty index reveals the metabolic potential of rare actinobacteria isolated from highly oligotrophic sediments. Microb Genomics 9:1–16.

29. Dinglasan JLN, Otani H, Doering DT, Udwary D, Mouncey NJ. 2025. Microbial secondary metabolites: advancements to accelerate discovery towards application. Nat Rev Microbiol 10.1038/s41579-024-01141-y.

30. Soldatou S, Eldjarn GH, Huerta-Uribe A, Rogers S, Duncan KR. 2019. Linking biosynthetic and chemical space to accelerate microbial secondary metabolite discovery. FEMS Microbiol Lett 366:1–8.

31. van der Meij A, Worsley SF, Hutchings MI, van Wezel GP. 2017. Chemical ecology of antibiotic production by actinomycetes. FEMS Microbiol Rev 41:392–416.

32. Fisch KM. 2013. Biosynthesis of natural products by microbial iterative hybrid PKS-NRPS. RSC Adv 3:18228–18247.

33. Wang H, Fewer DP, Holm L, Rouhiainen L, Sivonen K. 2014. Atlas of nonribosomal peptide and polyketide biosynthetic pathways reveals common occurrence of nonmodular enzymes. Proc Natl Acad Sci U S A 111:9259–9264.

34. Tyc O, Song C, Dickschat JS, Vos M, Garbeva P. 2017. The Ecological Role of Volatile and Soluble Secondary Metabolites Produced by Soil Bacteria. Trends Microbiol 25:280–292.

35. van Bergeijk DA, Terlouw BR, Medema MH, van Wezel GP. 2020. Ecology and genomics of Actinobacteria: new concepts for natural product discovery. Nat Rev Microbiol 18:546–558.

36. Alexander B. Chase, Douglas Sweeney, Mitchell N. Muskat, Dulce G. Guillén-Matus PRJ. 2021. Vertical Inheritance Facilitates Interspecies Diversification in Biosynthetic Gene Clusters and Specialized Metabolites. MBio 12:1–15.

37. Letzel AC, Li J, Amos GCA, Millán-Aguiñaga N, Ginigini J, Abdelmohsen UR, Gaudêncio SP, Ziemert N, Moore BS, Jensen PR. 2017. Genomic insights into specialized metabolism in the marine actinomycete Salinispora. Environ Microbiol 19:3660–3673.

38. Shi YM, Hirschmann M, Shi YN, Ahmed S, Abebew D, Tobias NJ, Grün P, Crames JJ, Pöschel L, Kuttenlochner W, Richter C, Herrmann J, Müller R, Thanwisai A, Pidot SJ, Stinear TP, Groll M, Kim Y, Bode HB. 2022. Global analysis of biosynthetic gene clusters reveals conserved and unique natural products in entomopathogenic nematode-symbiotic bacteria. Nat Chem 14:701–712.

39. Czech L, Hermann L, Stöveken N, Richter AA, Höppner A, Smits SHJ, Heider J, Bremer E. 2018. Role of the extremolytes ectoine and hydroxyectoine as stress protectants and nutrients: Genetics, phylogenomics, biochemistry, and structural analysis. Genes (Basel) 9:1–58.

40. Avalos M, Garbeva P, Vader L, Van Wezel GP, Dickschat JS, Ulanova D. 2022. Biosynthesis, evolution and ecology of microbial terpenoids. Nat Prod Rep 39:249–272.

41. Schöner TA, Gassel S, Osawa A, Tobias NJ, Okuno Y, Sakakibara Y, Shindo K, Sandmann G, Bode HB. 2016. Aryl Polyenes, a Highly Abundant Class of Bacterial Natural Products, Are Functionally Related to Antioxidative Carotenoids. ChemBioChem 17:247–253.

42. Valasek A, Kiss ÍÉ, Fodor I, Kovács M, Urbán P, Jámbor É, Fekete C, Kerepesi I. 2016. Proteomic insight into the primycin fermentation process of Saccharomonospora azurea. Acta Biol Hung 67:424–430.

43. Yamanaka K, Reynolds KA, Kersten RD, Ryan KS, Gonzalez DJ, Nizet V, Dorrestein PC, Moore BS. 2014. Direct cloning and refactoring of a silent lipopeptide biosynthetic gene cluster yields the antibiotic taromycin A. Proc Natl Acad Sci U S A 111:1957–1962.

44. Song PP, Zhao J, Liu ZL, Duan YB, Hou YP, Zhao CQ, Wu M, Wei M, Wang NH, Lv Y, Han ZJ. 2017. Evaluation of antifungal activities and structure–activity relationships of coumarin derivatives. Pest Manag Sci 73:94–101.

45. Dozio D, Sacchi F, Pinto A, Dallavalle S, Annunziata F, Princiotto S. 2025. Natural Antifungal Alkaloids for Crop Protection: An Overview of the Latest Synthetic Approaches. Pharmaceuticals 18.

46. Sequin CJ, Appelhans SC, Heis MS, Torrent WA, Trossero JA, Catalán CAN, Sampietro DA, Aceñolaza PG. 2023. Antifungal and toxicological evaluation of the alkaloids fraction from Neltuma nigra leaves. Biocatal Agric Biotechnol 54.

